# Spatial expression of an mRNA encoding Tie2-agonist in the capillary endothelium of the lung prevents pulmonary vascular leakage

**DOI:** 10.1101/2022.10.12.511878

**Authors:** Katrin Radloff, Birgitt Gutbier, Charlotte Maeve Dunne, Hanieh Moradian, Marko Schwestka, Manfred Gossen, Katharina Ahrens, Laura Kneller, Yadong Wang, Akanksha Moga, Leonidas Gkionis, Oliver Keil, Volker Fehring, Daniel Tondera, Klaus Giese, Ansgar Santel, Jörg Kaufmann, Martin Witzenrath

**Affiliations:** Pantherna Therapeutics GmbH, 16761 Hennigsdorf, Germany; Department of Infectious Diseases and Respiratory Medicine, Charité-Universitätsmedizin Berlin, corporate member of Freie Universität Berlin and Humboldt-Universität zu Berlin, 10117, Berlin, Germany; Institute of Active Polymers, Helmholtz-Zentrum Hereon, 14513 Teltow, Germany; Berlin-Brandenburg Center for Regenerative Therapies (BCRT) Charité Campus Virchow Klinikum, 13353 Berlin, Germany

## Abstract

Angiopoietin ligands Ang1 and Ang2 and the Tie2 receptor tyrosine kinases form an endothelial signaling pathway regulating vascular homeostasis and controlling vessel permeability, inflammation and angiogenic responses. Whereas Ang1-mediated Tie2 activation reduces inflammation and endothelial permeability, its antagonist, Ang2 increases it. Increased plasma Ang2 levels are associated with poor outcomes in patients with acute lung injury (ALI), as well as in acute respiratory distress syndrome (ARDS).

In the study presented here we tested the effect of a novel synthetic, nucleoside-modified mRNA-76 encoding for a hyperactive Ang1 derived fusion protein (COMP-Ang1) on attenuating post-inflammation vascular leakage. COMP-Ang1 mRNA was formulated into a cationic lipid nanoparticle (cLNP) using an optimized mixture of three different lipids and a microfluidic mixing technology. After intravenous injection, the respective mRNA-loaded LNPs were found to be delivered predominantly to the endothelial cells of the lung, while sparing other vascular beds. Also, the specific multimeric folding of the COMP-Ang1 protein complex appeared to be pivotal for its activity in preventing vascular leakage and in restoring the alveolar-endothelial barrier function in the inflamed and injured pulmonary vasculature. The mode of action of mRNA-76, such as its activation of the Tie2 signal transduction pathway, was tested by pharmacological studies *in vitro* and *in vivo* by systemic administration in respective mouse models. mRNA-76 was found to prevent lung vascular leakage/lung edema as well as neutrophil infiltration in an LPS-challenging model.

## Introduction

Acute respiratory distress syndrome (ARDS) is a life-threatening medical condition of respiratory failure characterized by rapid onset of widespread inflammation in the lungs. ARDS affects approximately 200 000 patients each year in the United States, and mortality and morbidity continues to be significant, remaining between 34.9 % and 40 %, depending on severity (Meyer et al. 2021; Matthay et al. 2019), resulting in nearly 75 000 deaths annually. Despite great improvements in management and heightened research efforts, ARDS still accounts for 10 % of intensive care unit (ICU) admissions globally, corresponding to more than 3 million patients with ARDS annually. ARDS can be caused by a variety of pulmonary (e.g., pneumonia, aspiration) or non-pulmonary (e.g., sepsis, pancreatitis, trauma) insults, leading to the development of nonhydrostatic pulmonary edema (Fan et al. 2018). Clinically, the syndrome is characterized by an acute developing, diffuse inflammatory lung injury initiated by increased alveolar capillary permeability, eventually resulting in increased lung weight, hypoxia (Fan et al. 2018) and fluid and neutrophil accumulation in the alveoli, which cannot be explained by hydrostatic edema. Primary treatment involves mechanical ventilation with low tidal volume together with treatments directed at the underlying cause, such as administering of antibiotics and fluid conservative strategies (Keddissi et al. 2019). However, to date there is no medication available for treatment and to further reduce ARDS mortality in patients. Hence there is a substantial medical need for novel and effective treatment approaches for ARDS, particularly in view of the significantly increased patient numbers affected by this life-threatening medical condition since onset of the worldwide “Covid-19” pandemic.

In ARDS, an initial vascular endothelial and alveolar epithelial cell activation or insult triggers massive liberation of inflammatory mediators, including complement activation products, cytokines, chemokines, proteases, and oxidants (Dolinay et al. 2012; Aranda-Valderrama and Kaynar 2018) which subsequently cause lung edema, a flooding of the lungs, thus impairing oxygen and carbon dioxide gas exchange of the lung capillaries. Albeit the inflammatory mediators recruiting immune cells, it is exactly their overabundance which eventually leads to a disruption of the endothelial barrier (vascular leakage) and to damage within the epithelial barriers in the alveolus. The endothelial angiopoietin (Ang)-Tie2 growth factor receptor pathway regulates vascular permeability and pathological vascular remodeling (Saharinen et al. 2017b). Ang1 and Ang2 are both multimeric ligands for the receptor tyrosine kinase Tie2. In particular, the inactivation of Tie2 signaling due to an inflammatory increase in Ang2 secretion is regarded to be one of the major molecular mechanisms for the impairment of barrier function in the lung vasculature. Conversely, activation by Ang1 of Tie2 signaling restores endothelial barrier function and prevents further vascular leakage. This unique function of the Ang-Tie2 pathway in vascular stabilization renders this pathway an attractive and validated target in conditions such as ARDS, sepsis, macular edema, and cancer (Saharinen et al. 2017a).

Antibodies specific for Ang2 have been employed to prevent microvascular endothelial barrier dysfunction and to attenuate vascular remodeling and inflammation in a chronic lung infection model (Tabruyn et al. 2010). However, an Ang2 neutralization dependent restoration of basal Tie2 phosphorylation has not been shown so far. The second promising therapeutic approach to overcome Ang2 repression of the normal Tie2 signaling pathway has been to drive Tie2 activation to supra-physiological levels using recombinant Ang1 protein itself as well as derivatives thereof, such as COMP-Ang1 (Cho et al. 2004a) or CMP-Ang1 (Khan et al. 2021; Cho et al. 2004b). Additionally, either an unrelated agonist such as *Vasculotide* (Wu et al. 2015; Gutbier et al. 2017) or administration of cells expressing Ang1 (McCarter et al. 2007) were shown to be applicable and effective to that purpose. The approach to prevent and restore vascular barrier function by directly utilizing Ang1 proteins or derivatives has been hindered so far by the inherent predisposition of Ang1 to form higher order complexes for it to be catalytically active, since the Tie2 receptor only interact with highly oligomeric Ang1. Also, multimerization of Ang1 displays unfavorable pharmacokinetics. Highly oligomeric native Ang1 variants are, however, difficult to produce, purify and store in a stable and active form, and hence production in their multimeric form under GMP conditions has not been possible for clinical use so far. Recombinant COMP-Ang1 is a fusion protein comprised of the multimerization domain of the rat cartilage oligomeric matrix protein (COMP) combined with the fibrinogen-related domain of human Ang1 and has been described previously as a hyperactive Tie2 agonist.

Here, we describe the mRNA-based approach of directed expression and secretion of a hyperactive Ang1 derivative, hCOMP-hAng1 fusion protein, to transiently activate the Tie2 pathway *in vitro* and *in vivo*.

## Materials & Methods

### mRNA synthesis

Codon-optimized mRNAs encoding human Ang1, COMP-Ang1, CMP-Ang1, Tie2 and NanoLuc proteins were synthesized and purified by BioSpring (Germany). Each mRNA was initiated with a cap, followed by a 5⍰ untranslated region (UTR), a signal peptide sequence, an open-reading frame (ORF) encoding Ang1, COMP-Ang1, CMP-Ang1 and Tie2, a 3⍰ UTR and a polyadenylated tail. Uridine was globally replaced with 1-methylpseudouridine.

### Cell transfection *in vitro*

Hela cells were transfected with mRNAs using Lipofectamine MessengerMax (Invitrogen, cat. #LMRNA001) according to manufacturer’s protocol. Briefly, cells were seeded in 150 cm^2^ culture flasks (TPP, cat. #90151) at 50 % confluency in serum-free DMEM (Gibco, cat. #61965-026). 15 h later, cells were transfected for 2 h with indicated amounts of MessengerMax-formulated mRNAs according to manufacturer’s protocol. Transfection media was removed and replaced with fresh serum-free DMEM for indicated time points. After 24 h, the supernatant was removed (around 25 ml) and either concentrated 10-fold with Amicon 3 kDa spin columns (Merck, cat. #UFC900308) and brought to a final volume of 2 ml for subsequent Western blot analysis (Fig.1B, C) or directly used for Tie2 activation studies (Fig. 2A, B, Fig. 3B). Human pulmonary microvascular endothelial cells (HPMEC) (Promocell, cat. #C-12281) were seeded onto 6-well plates (Thermo Scientific, cat. #140675) and left for 3 days until 100 % confluency in full supplemented ECM MV2 cell media (Promocell, cat. #C-22022). 15 h before transfection was placed in serum-reduced ECM MV2 media (containing 10 % of originally cell growth supplement MV2). Cells were transfected with preformulated mRNA-LNP at indicated concentrations in serum-free ECM media for 2 h. After transfection, media was replaced with serum-reduced ECM MV2 and left for indicated time points.

**Figure 1:**
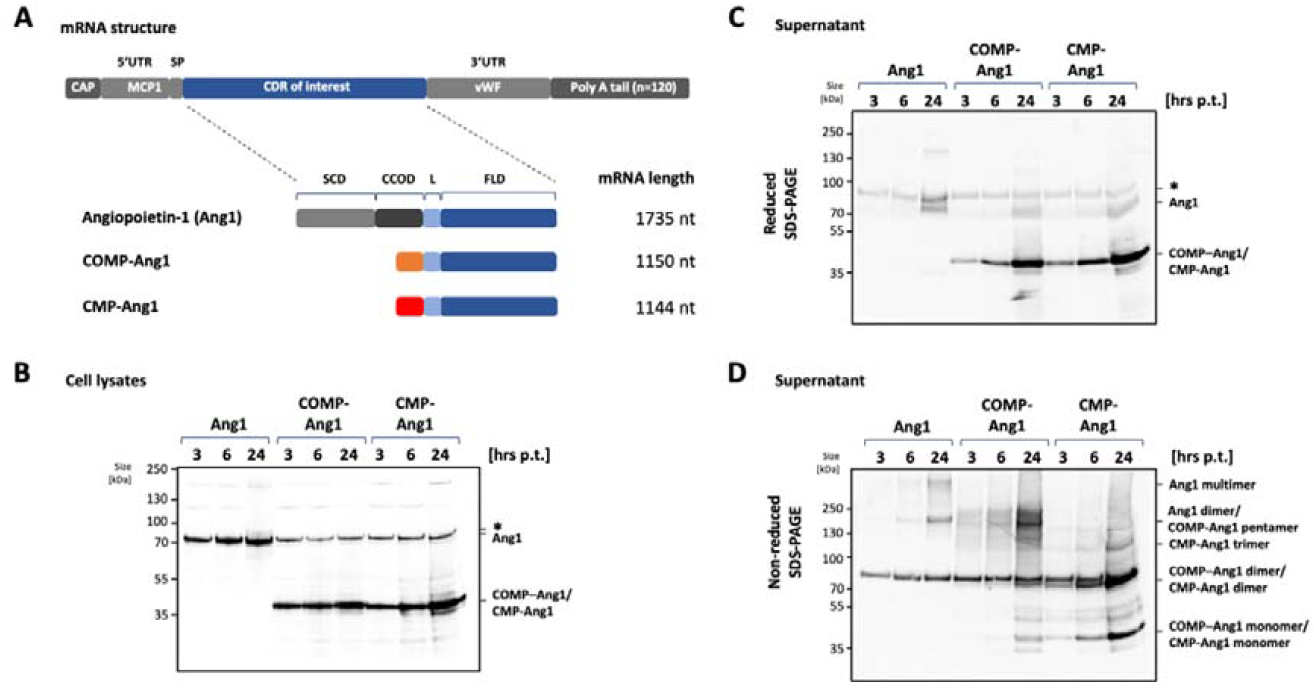
mRNA directed expression, secretion and multimerization of human Ang1, COMP-Ang-1 and CMP-Ang1 *in vitro*. **(A)** General structure of mRNA with signal peptide (SP) sequence flanked by designated 5’- and 3’-UTR regions plus defined polyA-tail. Schematic representation of protein domain structure of the natural protein Ang1 and the derived engineered humanized fusion proteins COMP-Ang1 and CMP-Ang1 according to Cho et al.,2004 and Oh et al., 2015. Protein domains of human Ang1: SCD, super clustering domain; CCOD, coiled coil domain; L, linker; FLD, Fibrinogen-related domain; the multimerization domains of COMP (orange) and of CMP protein (red) are indicated. **(B)** Expression, **(C)** secretion and **(D)** multimerization of wt Ang1 and Ang1 derivates in Hela cells. Cells were transfected for 2 h with mRNA-52 encoding Ang1, mRNA-76 encoding COMP-Ang1, or mRNA-59 encoding CMP-Ang1. Cell lysates and supernatants were harvested at indicated time points post transfection (p.t.) and analyzed by Western blotting using Ang1-specific antibody. (*, unidentified band)

**Figure 2:**
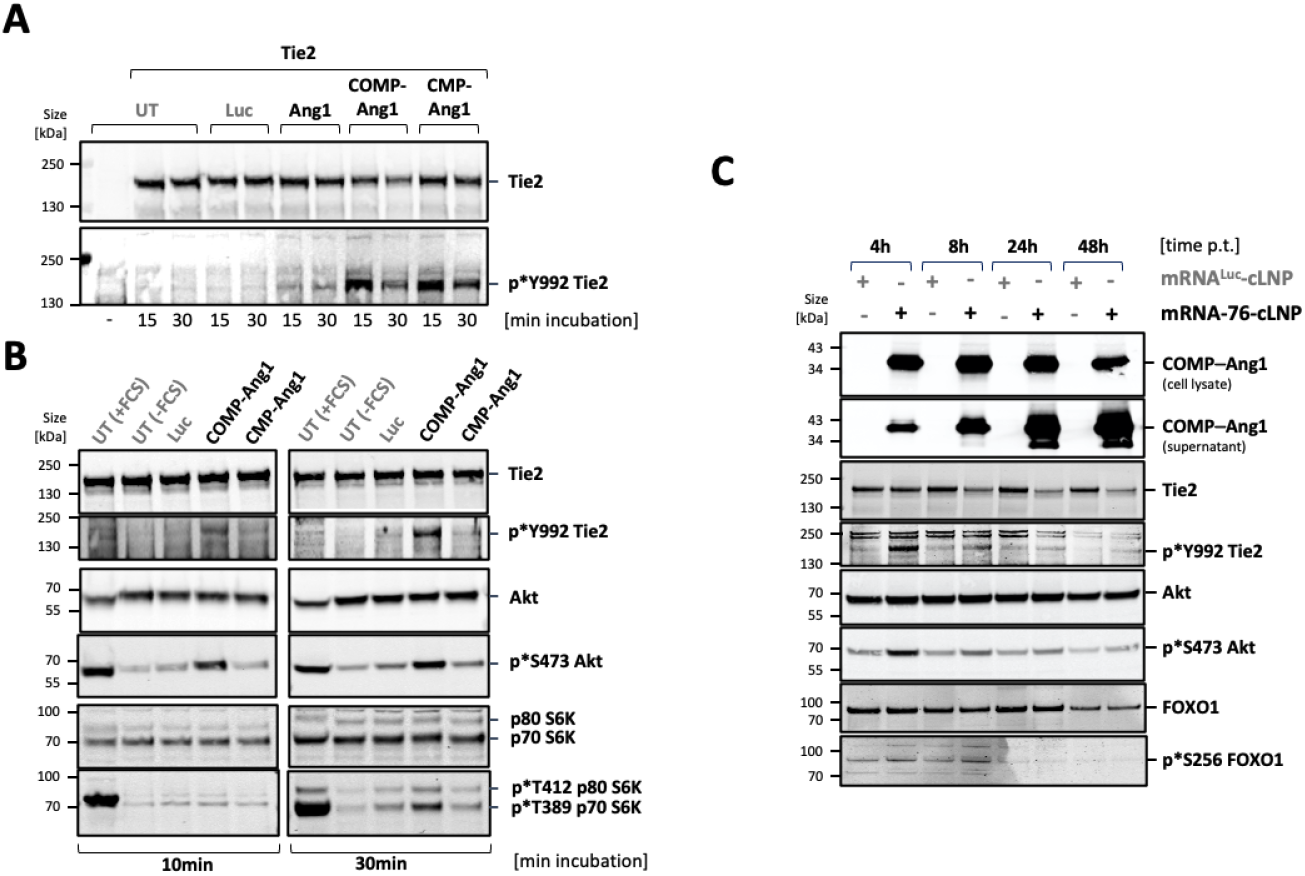
Activation of Tie2 pathway by Ang1, COMP-Ang1 and CMP-Ang1 containing supernatants. **(A)** Immunoblot of Tie2 expression and Tie2 phosphorylation from Tie2-expressing Hela cells after 15 or 30 min incubation with supernatant from HeLa cells expressing Luc, Ang1, COMP-Ang1 and CMP-Ang-1 (see Fig. 1) following incubation with supernatants from Hela cells transfected for 24h with Luc-mRNA or mRNA-52 encoding Ang1, mRNA-76 encoding COMP-Ang1, and mRNA-59 encoding CMP-Ang1. (B) Activation of Tie2 pathway in serum-starved primary human pulmonary microvascular endothelial cells (HPMEC) by COMP-Ang1 and by CMP-Ang1 containing supernatant from mRNA-76 or mRNA-59 transfected HeLa cells. Tie2-pathway activation is demonstrated by analysis of Tie2 phosphorylation status and by presence of phosphorylation at the downstream effectors Akt and S6K. **(C)** Tie2-pathway activation after transfection of LNP-formulated, COMP-Ang1-encoding mRNA-76 in serum-starved HPMEC. COMP-Ang1 expression, secretion, Tie2, Akt, and FOXO1 phosphorylation is shown by immunoblot on cell lysates at different time points post transfection (p.t.). (UT, untreated; FCS, Fetal calf serum; Luc, control Nano-Luciferase-encoding mRNA).

**Figure 3:**
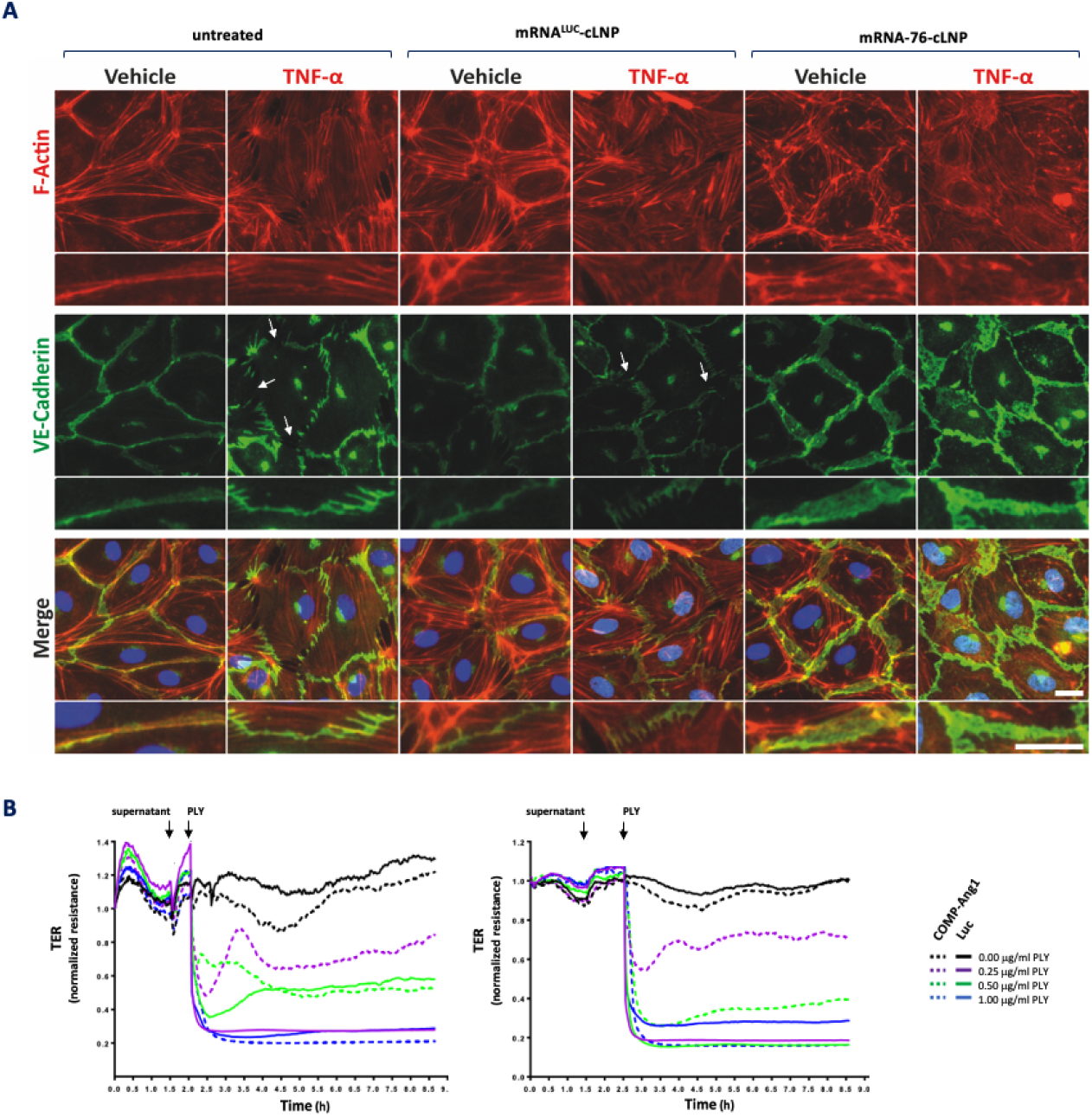
Cell staining by immunofluorescence demonstrating stabilization of endothelial adherens junctions. **(A)** Effect of COMP-Ang1 on F-actin remodeling and VE-Cadherin distribution in isolated human pulmonary microvascular endothelial cell. HPMECs grown on glass coverslips were transfected for 24h with control mRNA^LUC^ or with mRNA-76 encoding COMP-Ang1 with or without temporal addition of 10 ng/ml TNFα, followed by immunofluorescence staining with VE-Cadherin-specific antibody and with Alexa Fluor 594 phalloidin to detect actin filaments. Arrows indicate disrupted VE-Cadherin complexes. B) Transcellular electrical resistance (TER) measurements of HPMEC monolayers. HPMEC were incubated for 30 min (top) or 60 min (bottom) with supernatant from mRNA-76 or with mRNA^LUC^ (Luciferase) and the treated with supernatants from Hela cells transfected for 24 h with control mRNA^LUC^ or mRNA-76 encoding COMP-Ang1 and stimulated with indicated amount of pneumolysin (PLY; 0.25, 0.5, 0.75, 1.0 μg/ml). PLY stimulation decreased TER of HPMEC monolayers, displaying loss of endothelial integrity when incubated with control Luciferase supernatant (solid curve). Preincubation with COMP-Ang1 containing supernatant attenuated the PLY-induced TER decrease (purple lines dashed versus solid).

### Western Blot analysis

Cells were washed with ice-cold phosphate buffered saline (PBS) twice and lysed with ice-cold RIPA buffer (Thermo Scientific, cat. #89901) supplemented with Halt™ Protease and Phosphatase Inhibitor Cocktail (Thermo Scientific, cat. #78441). Tissue samples were solubilized in T-PER™ Tissue Protein Extraction Reagent (Thermo Scientific, cat. #78510) supplemented with Pierce Protease and Phosphatase Inhibitor tablets (Thermo Scientific, cat. #A32958) using Qiagen TissueLyser LT (Qiagen, cat. #69980). A mixture of cell or tissue lysate, 50 mM dithiothreitol NuPAGE reducing agent (Invitrogen, cat. #NP0009) and NuPAGE sample buffer (Invitrogen, cat. #NP0007) were heated at 70 °C for 10 min, electrophoresed in NuPAGE 4–12 % Novex Bis-Tris Gels and MOPS buffer (all from Invitrogen, cat. #NP0335BOX, NP0001) and transferred to nitrocellulose membrane (Cytiva, cat. #10600003; NuPAGE™ Transfer Buffer (Invitrogen, cat. #NP00061). After a 60 min blocking step (Intercept^®^ Blocking Buffer; LiCor, cat. #927-60001) membranes were washed with Tris Buffered Saline with Tween^®^ 20 (CST, cat. #9997S) probed at 4 °C overnight with specific primary antibodies diluted in Intercept Antibody Diluent (LiCor, cat. #927-65001). Binding of primary antibodies was detected by near infrared IRDye secondary antibodies (LiCor, cat. #926-32213, 926-32214, 926-68072) and near infrared signals were analyzed using LiCor Odysee CLx. Band intensities were analyzed using Empiria Studio Software 2.1.0134 (LiCor). Primary antibodies used: Angiopoeitin 1 (abcam, cat. #ab183701), human Tie2 (CST, cat. #7403), mouse Tie2 (Merck, cat. #05-584), pY992 Tie2 (R&D Systems, cat. #AF2720), Akt (CST, cat. #9272 and #2920S), pS473 Akt (CST, cat. #4060), S6K (CST, cat. #9202), pT389 S6K (CST, cat. #9234), FoxO1 (CST, cat. #2880), pT389 FoxO1 (CST, cat. #9461).

### Immunoprecipitation

Anti-Tie2 antibody ab33 (R&D Systems, Merck, cat. #05-584) for Tie2 IP or Tie2-Fc fusion protein (R&D Systems, cat. #313-TI) for COMP-Ang1 IP were covalently coupled to M-270 Epoxy Dynabeads™ (Thermo Scientific), cat. #14311D) at a ratio of 15 μg protein per mg Dynabeads according to manufacturer’s protocol. Briefly, 1.5 mg of antibody-coupled beads were incubated with tissue lysates overnight at 4 °C. Beads were washed 5 times with 1 ml with ice-cold RIPA buffer and finally denaturated using 1x NUPAGE loading dye containing 1x NUPAGE reducing agent (all Thermo Scientific). Beads were removed and lysates were analyzed by Western blotting analysis.

### Fluorescent microscopy

HPMEC were seeded in Chamber Slides (NUNCTM Lab-TekTM, cat. #154534PK) at 1×10^5^ cells/ml in endothelial MV2 growth medium. Cells were incubated for 3-5 days at 37°C in 5 % CO_2_. Cells were washed 2 times with pre-warmed PBS (Thermo Scientific, cat. #14190), final PBS wash was replaced with endothelial MV2 basal medium devoid of serum and supplements. Formulations containing mRNA were diluted in basal medium and added to each well for a final concentration of 0,38 ng/well. Cells were returned to the incubator for a further 2 hours. Media containing formulations was removed and replaced with 10 % MV2 growth medium for 16 h. For TNFα treatment, 10 % MV2 growth medium with TNFα (R&D Systems, cat. #210-TA) or vehicle was spiked into each well for a final concentration of 10 ng/ml for a further 6 h. Cells were fixed in 4 % PFA (Thermo Scientific, cat. #15670799) for 10 min at room temperature (RT). Cells were permeabilized in PBS containing 0.02 % Triton-X (PBS-T) for 2x 10 min washes on a rocker at RT. Primary antibody mix was made up with rabbit anti-VE-Cadherin (CST, cat. #2500) diluted at 1:400 in 2 % donkey immunobuffer containing 0.02 % Triton-X, rocking at 4 °C overnight. Primary antibodies were removed and washed in PBS-T 2x 10 min on a rocker at RT. Secondary antibody mix was made up with donkey anti-rabbit (H+L) highly cross-adsorbed Alexa FluorTM 647 (Thermo Scientific, cat. #A-31573) diluted at 1:400 and Alexa FluorTM 594 Phalloidin (Thermo Scientific, cat. #A12381) at 1:2000 in 2 % donkey immunobuffer containing 0.02 % Triton-X rocking at RT for 5 h. Cells were washed in PBS-T for a further 2x 10 min at RT. Nuclear staining was carried out with Hoechst (Tocris, cat. #5117) diluted at 1:500 in PBS for 15 min, followed by a final PBS-T wash. Fluorescent images were acquired with a Nikon Eclipse Ti-U with an ELWD S Pan Fluor 40X/0.6 objective lens. Images were processed using ImageJ (Schneider et al., 2012).

### Transcellular electrical resistance of human pulmonary microvascular endothelial cells

HeLa cells were first treated for 24 h with LNP-formulated mRNA-76 or with a Luciferase mRNA (see above) in order to generate conditional serum-free DMEM medium containing secreted COMP-Ang1 or secreted luciferase. Moreover, HPMEC were grown to confluency on evaporated gold microelectrodes (8-well array with ten electrodes per well, (ibidi GmbH, cat. #72010) and connected to an Electrical cell-substrate impedance sensing system (ECIS; Applied Biophysics, Troy, NY) (Tiruppathi et al. 1992; Giaever and Keese 1993) to enable continuous monitoring of transcellular electrical resistance (TER) (Mitchell et al. 1989). After monitoring baseline readings for 60-90 min, HPMEC medium was mixed 1:10 with the conditional HeLa media (COMP-Ang1 supernatant or Luc supernatant) for 30 min or for 60 min before treatment with three different concentrations of pneumolysin (PLY; 0.25, 0.5 and 1.0 μg/ml) (Mitchell et al. 1989) to evoke a barrier failure in HPMEC. TER values from each microelectrode were continuously monitored for 5 hours after PLY stimulation and normalized as the ratio of measured resistance to baseline resistance. The area under the curve (AUC) was calculated from start of PLY stimulation on.

### Animals

Female C57BL/6N mice (8 to 10 weeks, 18 to 20 g; Charles River, Sulzfeld, Germany) were used for all experiments. All animal experiments were approved by institutional and governmental German authorities (“Tierschutzbeauftragte” (animal welfare officer) and “Tierschutzausschuss” of Charité–Universitätsmedizin Berlin; and by the Landesamt für Gesundheit und Soziales Berlin; approval ID A0179/20), and were in accordance with the Federation of European Laboratory Animal Science Associations (FELASA) guidelines and recommendations for the care and use of laboratory animals (equivalent to American ARRIVE). Mice were randomly assigned to the appropriate experimental groups and were kept in closed, individually ventilated cages with filter hoods (type II-L, ZOONLAB), under specific pathogen-free conditions, with free access to food and water, room temperature between 20 and 22 °C, air humidity between 50 and 65 % and 12 h light/dark cycle.

### Transfection in vivo and isolated perfused and ventilated mouse lung (IPML) experiments

Intravenous tail vein administration (slow 100 μl bolus tail vein injection) with either Luciferase-mRNA (2 mg/kg) or mRNA-76 (2 mg/kg, 1.5 mg/kg or 1 mg/kg) was performed 6 h or 15 h before the experiment in the isolated mouse lung model was started. Subsequently, mice were anesthetized, placed in a 37 °C heated chamber (“artificial thorax”), tracheotomized and ventilated as described (Von Bethmann et al. 1998). After laparotomy, final blood collection via the vena cava, sternotomy, and cannulation (pulmonary artery, left atrium), lungs were perfused with Krebs-Henseleit-hydroxyethylamylopectine buffer (Serag-Wiessner) supplemented with sodium bicarbonate (1:50). The chamber was closed, and lungs were ventilated by negative pressure (Pexp -4.5, Pins -9.0 cm H_2_O) and perfused (1 ml/min) for 20 min to establish baseline conditions. Then 0.04 % human serum albumin (HSA; 20 % solution; CSL Behring, cat. #Human-Albumin 20 % Behring) was admixed continuously to the perfusate 10 min prior to intravenous bolus application of recombinant pneumolysin (PLY; 1.4 μg/ml) (Mitchell et al. 1989). Thirty minutes after PLY challenge, bronchoalveolar lavage (BAL) was performed (2 x 650 μl, 0.9 % saline buffer), and HSA concentration was measured in BAL fluid (BALF) via ELISA (Bethyl Laboratories, cat. #E88-129) in order to quantify the lung barrier failure (Seybold et al. 2005; Witzenrath et al. 2006).

### Neutrophilia transmigration experiment

Animals (n=10) were challenged intratracheally with 0.9 % w/v saline or with LPS (3 mg/kg). Two hours later animals were treated with mRNA-76-cLNP or with control mRNA^LUC^-cLNP employing a dosage of 1.5 mg/kg via intravenous tail vein injection. Control group animals received intraperitoneal injections of Dexamethasone (3 mg/kg) one hour prior to, and 8 hours after LPS administration. Animals were sacrified after 24 hours and the airway lavaged with 0.3 ml of PBS using an inserted cannula. The bronchoalveolar lavage from each mouse was centrifuged at 1000 g for 10 min at ca 4 °C. The cell pellet was re-suspended in 0.5 ml PBS and retained on wet ice until analysis. A total and differential cell count of the BALF supernatant was performed using the XT-2000iV (Sysmex UK Ltd). The samples were vortexed for approximately 5 seconds and analyzed. Total and differential cell counts (including neutrophils, lymphocytes and mononuclear cells (includes monocytes and macrophages)) were measured as number of cells per animal.

## Results

Expression of coding sequences of the wild-type human Ang1 and of engineered fusion proteins COMP-Ang1 and CMP-Ang1 flanked by heterologous untranslated regulatory sequences was tested in cell culture systems in order to identify a potent mRNA construct for Tie2 activation. Wildtype human Ang1 mRNA with heterologous UTRs served as a control next to either a construct with an ORF for a COMP-Ang1 fusion protein or a construct containing an ORF with a CMP-Ang1 fusion protein **(Fig. 1A).** Similar COMP-Ang1 and CMP-Ang1 recombinant fusion proteins have been previously described which contain the rat homologue of COMP or human CMP as potent Tie2 agonist in *in vitro* assays and in mouse models (Cho et al. 2004b; Oh et al. 2015; Wallace et al. 2021).

### mRNA directs transient expression of human Ang-1 and COMP-Ang1 and CMP-fusion proteins *in vitro*

*mRNAs* encoding wildtype human Ang1, fusion proteins COMP-Ang1 and CMP-Ang1 were transfected into HeLa cells using Lipofectamine MessengerMax. Cell culture supernatants were collected 3 h, 6 h and 24 h after the transfection, and corresponding whole cell lysates were prepared. The supernatants and cell lysates were subsequently analyzed by immunoblotting employing an Ang-1 specific antibody. A strong protein expression signal was detected for all three constructs, and protein levels post-transfection of the mRNA construct were found to be higher than those of endogenous Ang-1 protein **(Fig. 1B).** Equivalent protein levels were detected in the supernatant of the cell culture, indicating secretion of the expressed proteins **(Fig. 1C).** Multimerization potential of the expressed Ang1 derivates was analyzed by employing non-reducing PAGE on the supernatant. Contrary to CMP-Ang1, the COMP-Ang1 fusion protein was found to be capable of forming higher-order (pentameric) homopolymers **(Fig. 1D).**

### Activation of Tie2 by COMP-Ang1 and CMP-Ang1

The chimeric COMP-Ang1 derivative encoded by mRNA-76 was found to *in vitro* activate Tie2 in a paracrine manner. Tie2-negative HeLa cells were first transfected for 24 h with *mRNA* encoding for Tie2 receptor using Lipofectamine MessengerMax. Subsequently, cells were incubated for 15 min or 30 min with supernatants from HeLa cells previously transfected for 24 h with mRNA encoding the different Ang1 derivatives, or with NanoLuc-encoding control mRNA using Lipofectamine MessengerMax. Ensuing, Tie2-transfected HeLa cells were lysed and protein levels analyzed by Western blotting. Both supernatants with Ang1 derivatives displayed a significantly stronger activation of the Tie2 receptor as assessed by phospho-Y992 Tie2 antibody **(Fig. 2A)** than when activated by the Luc-control and wildtype Ang1 proteins. Ability of different Ang1-derivatives to induce paracrine activation of the Tie2 pathway in human primary endothelial cells was then investigated in a second set of experiments. For this, starved primary HPMECs (human primary microvascular endothelial cells) were incubated for 10 and for 30 min with supernatants from HeLa cells previously transfected in serum-free medium for 24 h with mRNAs encoding the different Ang1 derivatives using Lipofectamine MessengerMax, or with NanoLuc as a control. HPMEC cells were lysed after 10 and 30 min, and phosphorylation status of Tie2, Akt and S6K analyzed by Western blotting and compared to lysates from starved cells or from cells stimulated for 10 and 30 min with FCS **(Fig. 2B).** Strongest downstream activation of Akt and S6K was seen with FCS treatment on starved cells, thus corroborating the expected broad growth factor response. Cells treated with supernatant from HeLa cells transfected with mRNA-76 encoding COMP-Ang1 also showed robust phosphorylation of Akt and of S6K, but additionally exhibited autophosphorylation of Tie2, indicative for a more specific pathway activation. The paracrine effect of HeLa conditional medium-derived COMP-Ang1 resulted in a better Tie2 pathway activation as compared to CMP-Ang1, leading to selection of mRNA-76 /COMP-Ang1 for the following experiments. mRNA-76 encoding COMP-Ang1 was directly transfected into HPMEC and secretion and Tie2 signaling was analyzed in order to demonstrate increased autocrine Tie2 activation. Strong COMP-Ang1 expression was observed in cell lysates and in the supernatant of the respective culture, indicating efficient secretion **(Fig. 2C).** Lysate analysis of serum-starved HPMEC transfected with mRNA-76 revealed phosphorylation of Tie2, Akt and FOXO1, thereby indicating functional Tie2 pathway activation at 4 h and 8 h post-transfection, as well as fast onset of activation. Waning of Tie2 receptor activation and a reduction of Tie2 receptor protein was observed at 24 h and 48 h post transfection, consistent with previous reports describing a ligand dependent Tie2 internalization and degradation (Bogdanovic et al. 2006; Ghosh et al. 2016; Wehrle et al. 2009) as well as reduced expression of Tie2 mRNA upon Ang1 treatment (Hashimoto et al. 2004).

### COMP-Ang1 expression stabilizes endothelial barrier function *in vitro*

Regulation of endothelial barrier stabilization by Ang1/Tie2 signaling has been shown to be mediated by VE-Cadherin-dependent cell-cell adhesion and cortical actin formation, since the cytoplasmatic tail of VE-cadherin is known to contain several phosphorylation sites with different distinctive and selective effects on endothelial cell function (Hellenthal et al. 2022). However, inflammatory conditions lead to disruption of VE-Cadherin complexes by phosphorylation-dependent internalization and by degradation, as well as to formation of actin stress fibers. We analyzed VE-Cadherin distribution in HPMEC cells upon TNFα challenge by immunofluorescence staining and by microscopy analysis.

As reported previously (Xia et al. 1998), TNFα was shown to induce disruption of cell-cell adhesion as verified by disruption intercellular VE-Cadherin staining **(Fig. 3A,** arrows) and by formation of actin stress fibers in non-transfected cells (UT) as well as in cells transfected with control mRNA^LUC^-cLNP. However, cells transfected with mRNA-76-cLNP displayed more VE-Cadherin staining even if unstimulated but in the presence of TNFα. This observation argues for the existence of increased VE-Cadherin trans-interaction area at overlapping cell edges, as previously also described by Birkuva *et al*. (Birukova et al. 2012).

Following demonstration of functional Tie2 activation on VE-Cadherin complex stabilization, we investigated the potential effect of mRNA-76 directed expression of COMP-Ang1 on pneumolysin-evoked barrier failure in cell culture. Recovery of the pneumolysin-evoked barrier failure was measured by monitoring transcellular electrical resistance (TER) (Srinivasan et al. 2015). TER measurements are performed by applying an AC electrical signal across electrodes placed on both sides of a cellular monolayer and subsequently measuring voltage and current to calculate the electrical resistance of the barrier (Elbrecht et al. 2016). A significantly improved transcellular electrical resistance was observed with the mRNA-76 conditional medium at lowest pneumolysin concentration of 0.25 μg/ml, but not with the supernatant form HeLa cells transfected with Luciferase mRNA **(Fig. 3B,** purple lines: COMP-Ang1 dotted line, Luciferase supernatant solid line). This effect was observed in pneumolysin-induced barrier failure experiments after pretreatment with conditional supernatant for 30 min or 60 min. These data indicate that mRNA-76 treatment leads to expression and secretion of COMP-Ang1 and the so expressed COMP-Ang1 is functional in a paracrine manner on microvascular endothelial cell (HPMEC) monolayers as demonstrated by its ability to attenuate the PLY-induced TER decrease. These findings confirm the proposed mode of action of our COMP-Ang1 molecule in diminishing inflammation induced hyperpermeability via its respective activation of the Tie2 signaling pathway.

### mRNA-76 enables COMP-Ang1 expression and Tie2 pathway activation in mice

Various physical characteristics of LNPs affect organ biodistribution and cell specificity of the mRNA delivery, namely size, surface charge, pKa of the lipids, lipid composition and mRNA/lipid ratio, PEGylation (Eygeris et al. 2022; Hou et al. 2021; Kauffman et al. 2016; Kowalski et al. 2019). We show that our mRNA-76 formulated in cationic lipid-nanoparticles (cLNPs) enable Tie2 activation by paracrine and autocrine COMP-Ang1 expression *in vivo*.Our cationic LNP formulations are based on lipoplex formulations previously described, for siRNA *in vivo* delivery to the pulmonary endothelium (Fehring et al. 2014; Gutbier et al. 2018; Santel et al. 2006b, 2006a). Expression of mRNA-76 directed COMP-Ang1 was analyzed by Western Blot of protein from lung tissue from mice (n=4) treated by intravenous tail vein administration with 2 mg/kg mRNA-76-cLNP. Robust COMP-Ang1 expression was detected in lung lysates from mice 6 h after treatment, and declining expression levels were observed in mice 24 h post treatment due to the transient nature of mRNA **(Fig. 4A).** As expected from previous studies with cationic LNPs, mRNA-76-cLNP mediated robust lung tissue-selective COMP-Ang1 expression, while inducing no significant COMP-Ang1 expression in tissue lysates from heart, liver, spleen, and kidney **(Fig. 4B).** Immune-precipitation was performed on different organ lysates using Tie2-Fc fusion protein for capturing of COMP-Ang1 to confirm lung selective COMP-Ang1 expression. Even with this more sensitive detection method the predominant COMP-Ang1 expression was observed in lung lysates, with some minor expression also seen in the spleen **(Fig. 4C).** No significant expression of COMP-Ang1 was detected in liver, heart and kidney thus confirming the selective pulmonary delivery of mRNA-76-cLNP.

**Figure 4.**
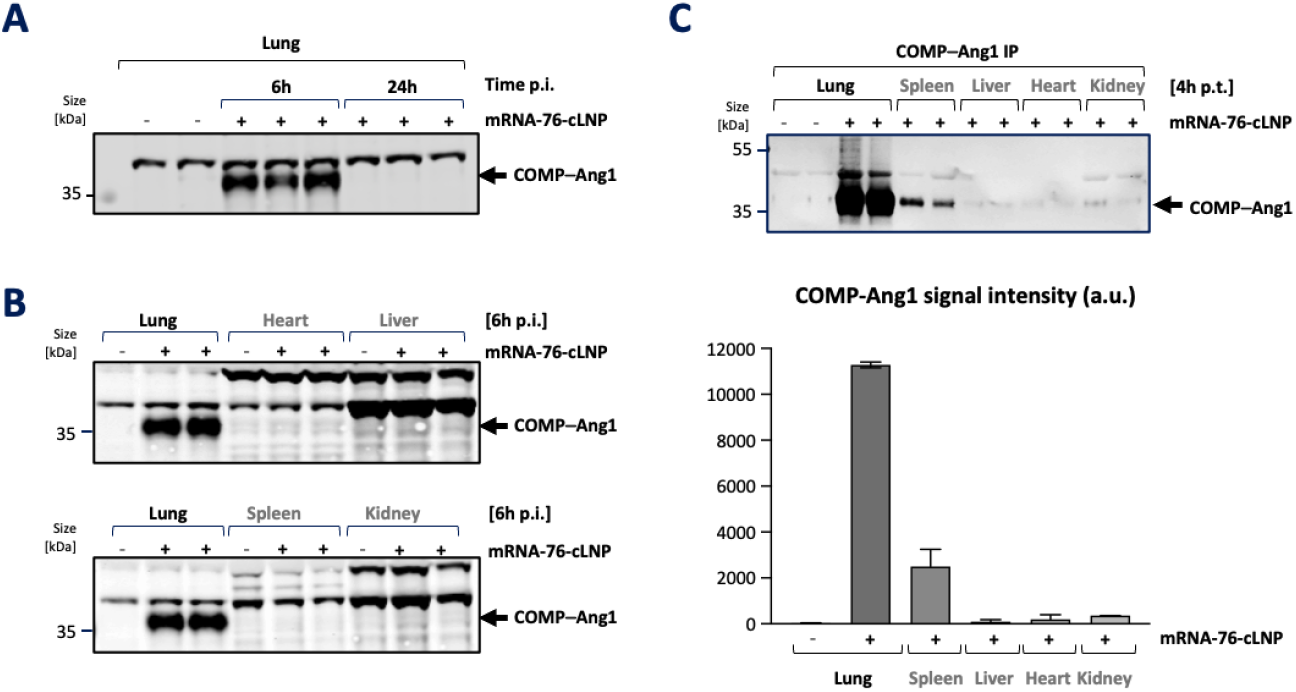
Transient expression of COMP-Ang1 in murine lungs by cLNP-mediated lung delivery of mRNA-76. **(A)** Immunoblot of lung tissue lysates 6 and 24 h after administration of 1.5 mg/kg mRNA-76-cLNP (+) or saline control (-). The expressed COMP-Ang1 fusion protein is indicated by an arrow. **(B)** Immunoblot of different tissue lysates 6h post-administration of 1.5 mg/kg mRNA-76-cLNP (+) or saline control (-). Lung selective COMP-Ang1 protein is indicated by an arrow. **(C)** Immunoprecipitation of COMP-Ang1 protein using Tie2-Fc-coupled magnetic beads from indicated tissue samples 4 h after administration of 1.5 mg mRNA-76-cLNP (+) or saline control (-).

### Single-cell transcriptomics of murine lungs after mRNA-76-cLNP treatment

We performed single-cell RNA-sequencing on cells derived from murine lungs in order to characterize the specific cell types with mRNA-76 uptake. Lungs from two mice treated with 1.5 mg/kg mRNA-76-cLNP (2 h post administration) were obtained, and lung cell suspension was subjected to droplet-based single-cell gene expression profiling **(Fig. 5A).** We obtained 5,100 high-quality transcriptome readings and were able to identify 13 cell clusters with main cell types of the respective lung samples by employing dimensionality reduction via t-distributed stochastic neighbor embedding (t-SNE) and graph-based clustering **(Fig. 5B).**

**Figure 5:**
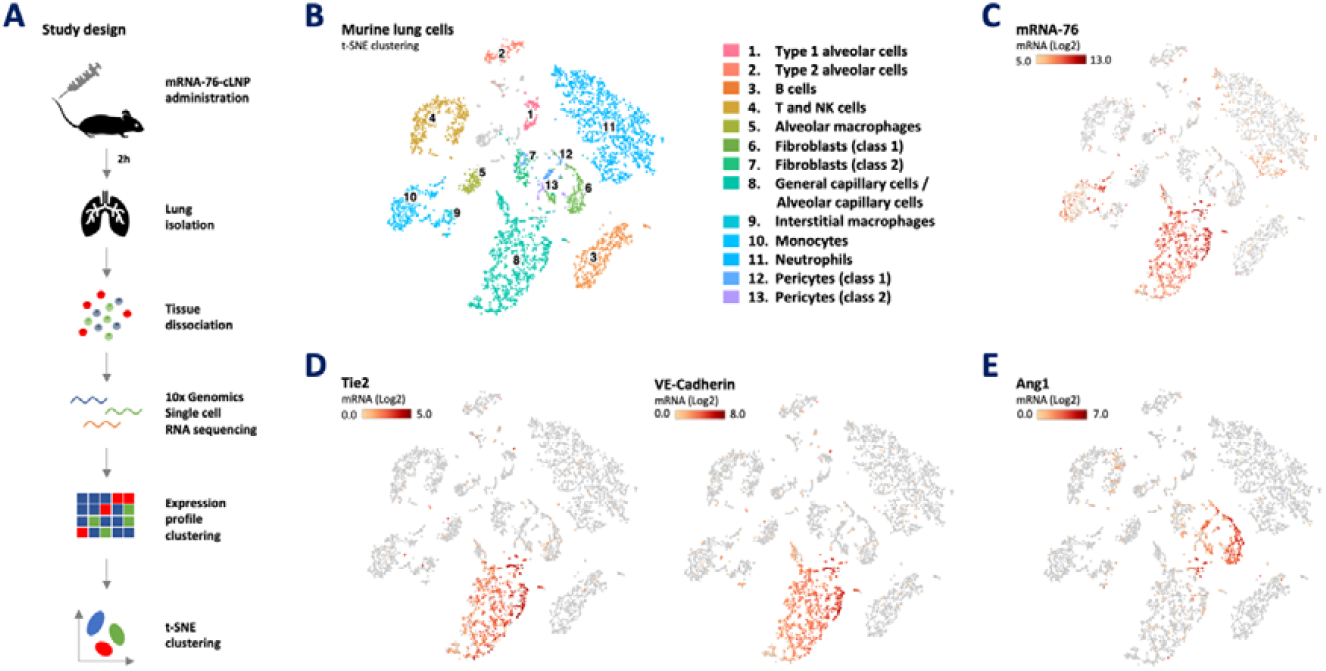
cLNP-mediated delivery of 1.5 mg/kg mRNA-76 leads to uptake predominantly in murine pulmonary capillary endothelial cells. **(A)** Experimental design and workflow **(B)** Loupe-based t-distributed stochastic neighbor embedding (t-SNE) clustering of murine lung cell types clustered by single-cell transcriptional analysis of 5,100 cells after quality control, showing a total of 13 distinct cell type clusters as indicated by numbers. **(C)** t-SNE plots showing levels for COMP-Ang1-encoding mRNA-76. **(D)** t-SNE plots showing levels for canonical markers of endothelial cells (Tie2, VE-Cadherin). **(F)** cells expressing Ang1 (pericytes class 1 and 2, fibroblasts class 1). The intensity of expression is indicated by red coloring.

We observed mRNA-76 reads to be largely restricted to cluster 8 and cluster 10, representing cells of the alveolar capillary endothelium (gCap, general capillary cells) and aCap (alveolar capillary cells) recently termed aerocytes (Gillich et al. 2020), as well as monocytes, respectively. Minor amounts of mRNA-76 appeared to also be present in a sub-cluster of the pulmonary neutrophils (cluster 11). We analyzed the mRNA reads of Tie2 and VE-cadherin, both being endothelial cell specific expressed genes, in order to confirm the endothelial specific clustering for mRNA-76 into cluster 8. The overlap with the reads for both markers confirmed cell type specificity of mRNA-76 delivery **(Fig. 5D).** In contrast to mRNA-76, endogenous Ang1 mRNA is mainly detected in cluster 6, which does not overlap with reads for the Tie2 receptor mRNA **(Fig. 5E).** The observed cell type-specific expression profiles suggest a paracrine Tie2 activation for the endogenous Ang1, as well as a potential autocrine Tie2 activation by ectopically expressed, mRNA-76-encoded COMP-Ang1.

### mRNA-76 mediated Tie2 pathway activation in murine lungs

We then demonstrated Tie2 pathway activation in lung tissue to be induced by mRNA-76 COMP-Ang1 expression in murine lung tissue. Activation of PI3 kinase signaling including Akt phosphorylation is known to be a downstream effect of Tie2 receptor activation. Of note, demonstrating COMP-Ang1 mediated activation of Tie2 signaling is difficult due to the low proportion of Tie2-expressing endothelial cells within total lung lysates in healthy, non-inflamed lungs. Therefore, Tie2 was immuno-precipitated from lung tissue lysates of mice previously treated with mRNA-76 using Tie2-specific antibodies. Tie2 phosphorylation of the lung was subsequently shown by Western blot analysis of immuno-precipitated samples using anti-phospho-Tie2 antibody (p*Y992) **(Fig. 6A).** Signal quantification of phosphorylated Tie2 was performed using LiCor Odyssee CLx and measured relative to corresponding total Tie2 levels. In addition, immunoblot analysis using Ang1-specific antibodies showed co-immunoprecipitation of COMP-Ang1 thus demonstrating efficient binding of human COMP-Ang1 to murine Tie2. Since Akt phosphorylation upon COMP-Ang1 stimulation is mediated by Tie2 activation, whole lung tissue lysates from mice treated with 1.5 mg/kg mRNA-76-cLNP were analyzed for COMP-Ang-1 expression, Akt expression and Akt-phosphorylation by immunoblot **(Fig. 6B)** in comparison to control lysates (Saline treated). Akt phosphorylation levels were shown to be highest 6 h post mRNA-76-cLNP treatment.

**Figure 6:**
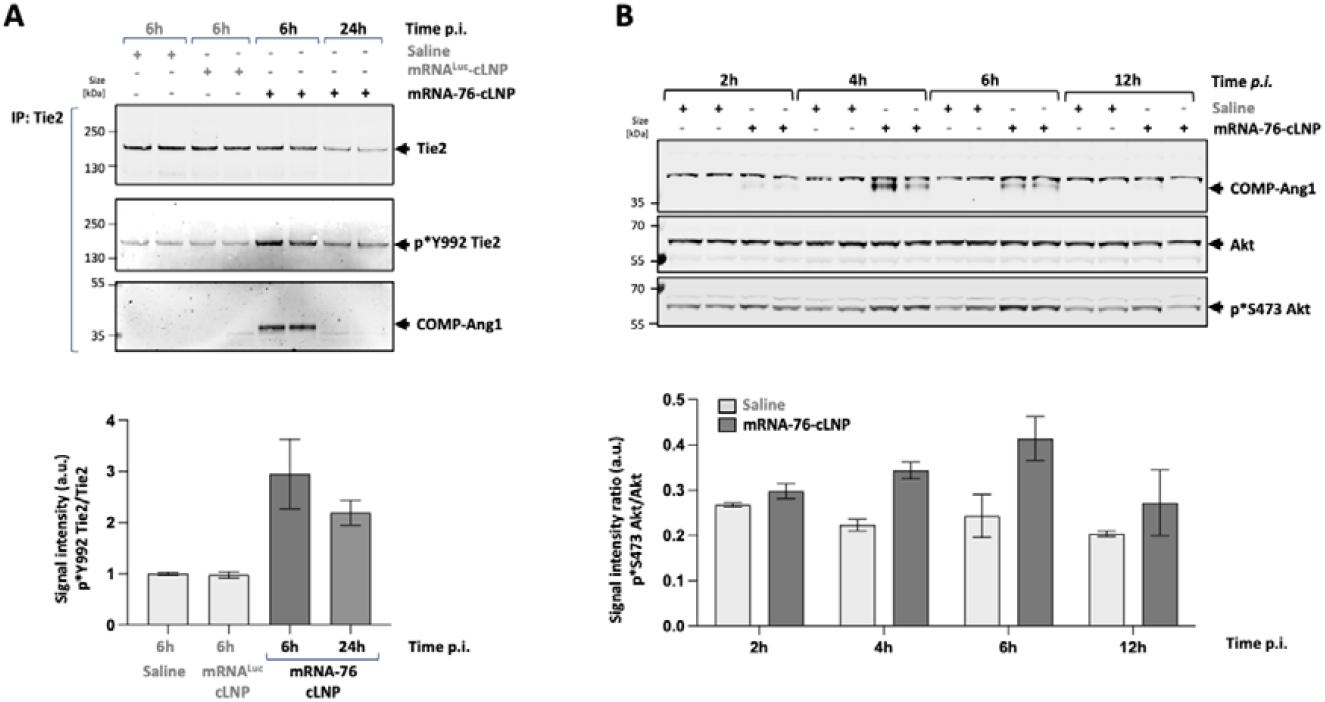
cLNP-mediated delivery of mRNA-76 leads to time-dependent Tie2 pathway activation *in vivo*. **(A)** Immunoblot after immunoprecipitation of murine Tie2 from lung samples at indicated time points after i.v. administration of 1.5 mg/kg mRNA-76-cLNP. **(B)** Immunoblot of murine lunge tissue lysates analyzing COMP-Ang1 expression level and Akt phosphorylation status at indicated time points after i.v. administration of 1.5 mg/kg mRNA-76-cLNP or Saline.

### mRNA-76 treatment reduces pneumolysin-evoked vascular permeability in an isolated perfused and ventilated mouse lung (IPML) model for vascular leakage

A pneumolysin (PLY) stimulation experiment was performed on isolated and ventilated mouse lungs (IPML model) to demonstrate mRNA-76 efficacy in preventing and/or decreasing lung hyperpermeability by inflammation **(Fig. 7A).** In the employed IPML model, PLY stimulation increases lung permeability 30 min after application, as described previously (Gutbier et al. 2017; Witzenrath et al. 2006). Here, treatment with mRNA-76 significantly decreased pneumolysin-induced hyperpermeability of mouse lungs as compared to treatment with Luciferase **(Fig. 7B).** Dose dependent reduction of HSA levels in the BALF was observed at 6 h and 15 h post mRNA-76 treatment as compared to that derived from control mice, indicating reduced vascular leakage. These results suggest that mRNA-76 pretreatment is effective in preventing pneumolysin-induced vascular leakage in the IPML model, and respective observations are consistent with the previously established mode of action of Tie2 activation by mRNA-76 directed expression of COMP-Ang1 in lung tissues. The effective dose in this mouse efficacy model is approx. 1 mg/kg for the 6 h time point and marginally higher at later time points (15 h). This decreasing effect on vascular leakage at later time points with lower doses suggests a more sustained expression with higher mRNA-76 dosing.

**Figure 7:**
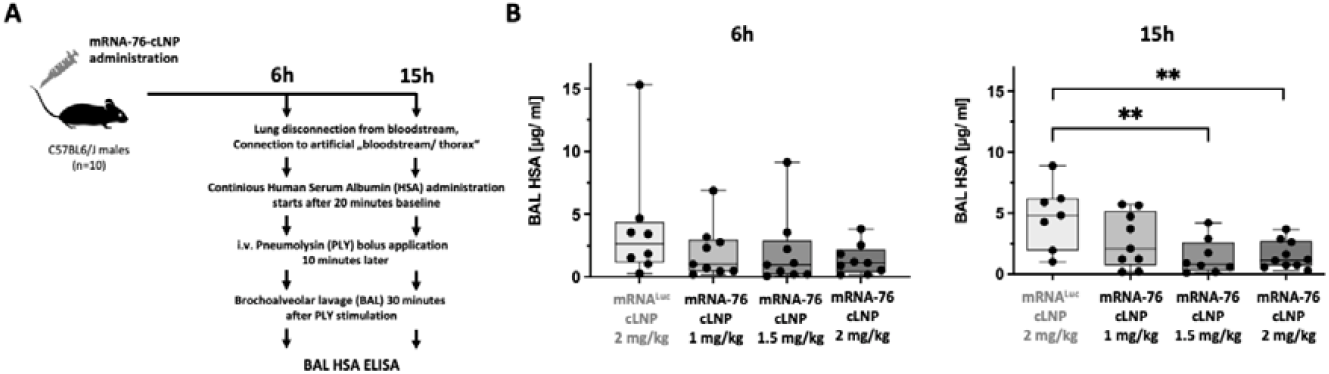
Systemic mRNA-76 treatment stabilized endothelial barrier lung function in an *ex vivo* isolated perfused and ventilated mouse lung (IPML) model. **(A)** Schematic diagram of the experimental protocol of the *ex vivo* perfused and ventilated mouse lung model. 6 h, 15 h after administration of mRNA-76-cLNP (1, 1.5 and 2 mg/kg) or control mRNA^LUC^ -LNP (2 mg/kg) lungs were isolated and stimulated with PLY (1.4 μg/ml) for 1 minute. After 30 min, lung vascular permeability was assessed by quantifying respective concentrations of continuously infused human serum albumin (HSA) in the bronchoalveolar lavage fluid (BALF). **(B)** Graphic showing treatment with mRNA-76 to significantly decrease pneumolysin-induced hyperpermeability of mouse lungs as compared to treatment with control mRNA^LUC^-LNP 6 h (left) and 15 h (right) post mRNA-76-cLNP treatment as shown by HAS ELISA. Values are given as mean (n⍰=⍰10, **p<0.01 between indicated groups) concentration of human serum albumin (HSA) in bronchoalveolar lavage fluid (BALF). HSA concentration of individual mice are indicated as dots.

### mRNA-76 directed COMP-Ang1 expression reduces neutrophil influx after LPS challenge in murine lungs

We observed a significant therapeutic effect on endotoxin/LPS induced neutrophilia after temporal and spatial expression of COMP-Ang1 in a murine model of endotoxin (*E. coli* lipopolysaccharide, LPS)-induced pulmonary inflammation **(Fig. 8A).** Hence, we detected a significant decrease in LPS induced neutrophil influx into the alveoli after single mRNA-76-cLNP treatment in our experimental model **(Fig. 8B).** The level of the decreased neutrophil influx effect is in the same range as that of prophylactic and therapeutic dexamethasone treatment used as positive control for this assay. No significant reduction regarding cell influx was observed with mRNA^Luc^-cLNP as compared to vehicle control mice. Also, no changes in LPS induced vascular leakage (e.g., changes in wet lung weights) were observed with mRNA-76-cLNP treatment as compared to control mRNA formulation, indicating that vascular leakage and leucocyte emigration do not necessarily occur together in the blood vessels of the lung. The fact that elevated levels of circulating cytokines are not perturbed by enhancement of vascular integrity and hence the notion that leakage can be regulated distinctly from the inflammatory response has been discussed previously by others (Filewod and Lee 2019). Generally, the effect of mRNA-76-cLNP treatment in rodent models for LPS-induced lung injury could also be dependent on the time point at which samples are obtained, and at which respective physiological measurements are taken. The LPS model might therefore be not adequate to simultaneously address the question of both neutrophil efflux as well as lung edema at similar timepoints 24 h post endotoxin challenge, as opposed to the IPML model which was used primarily for experimentally reducing vascular leakage. In our in a murine model of LPS-induced pulmonary inflammation, intravenous mRNA-76-cLNP treatment showed a strong therapeutic effect in reducing leukocyte extravasation and/or transmigration of neutrophils for the timepoint(s) analysed. Notably, leukocyte influx reduction and reduction in edema formation might be clinically highly desirable to counteract delayed apoptosis of parenchymal neutrophils, believed to contribute to tissue injury (Teder et al. 2002). Taken together, our experiments in the IPML mouse model confirm a dose dependent therapeutic effect of COMP-Ang1 on endothelial permeability, as well as a therapeutic efficacy on dysregulated lung inflammation in the LPS-induced pulmonary neutrophilia mouse model. Both underlying pathophysiological mechanism, loss of endothelial permeability and dysregulated lung inflammation, are hallmark events of early ARDS syndrome.

**Figure 8:**
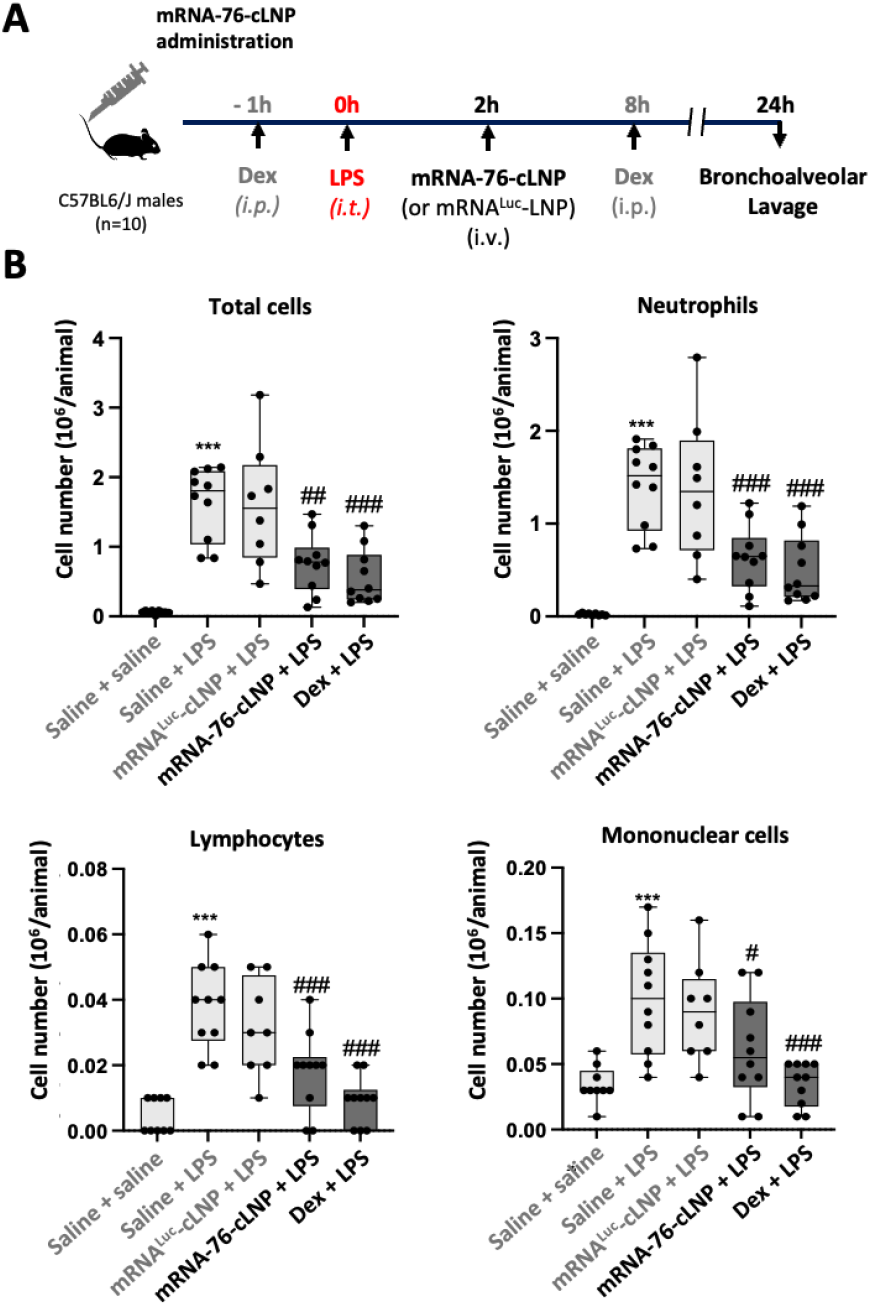
LPS-induced Pulmonary Neutrophilia Mouse Model. **(A)** Schematic drawing of the experimental protocol. Animals (n=10) were challenged intratracheally with 0.9% w/v saline or LPS (3 mg/kg). A fixed volume of 50 μL, equal to an approximate intratracheal dose volume of 2.5 mL/kg, based on a 20 g mouse, was applied intratracheally. 2h after LPS administration mRNA-76-cLNP (1.5 mg/kg IV) or mRNA^LUC^-cLNP01 (1.5 mg/kg IV) was intravenously administered, and bronchoalveolar lavage fluid (BALF) was analysed after 24 h. **(B)** Effect of COMP-Ang1 expression on BALF total and differential cell counts and on wet lung weight in a murine model of endotoxin (LPS)-induced pulmonary inflammation. Data shown as mean ± standard error of the mean. *** p<0.001 when compared to saline challenged control group. #p<0.05, ##p<0.01, ###p<0.001 as compared to LPS challenged and vehicle treated group.

## Discussion

In our work presented here we introduce mRNA-76-LNP, a novel drug modality utilizing mRNA-based expression technology, in combination with a lipid nanoparticle-based delivery technology, to achieve transient and localized expression of functional Ang1 protein derivatives in the lung vasculature. Our new approach and technology thus overcome the limitations of efficaciously administering recombinant multimeric proteins for activating Tie2 signaling. For example, full-length Ang1 unlike Ang2, has been shown previously by Xu et al. to be incorporated efficiently into the extracellular matrix (ECM) via its linker peptide region, thereby explaining its rapid serum clearance. Interestingly, in our experiments, only COMP-Ang1 and CMP-Ang1 were detected in significant amounts in the cell supernatant, thus indicating efficient secretion **(Fig. 1C).** This is in contrast to the comparably high intracellular expression levels in lysates of HeLa cells transfected with the three Ang1 variants **(Fig. 1B).** One possible explanation for this observed difference might be that albeit the Ang1 linker domain being present in all three Ang1 variants, conceivably only wt-Ang1 interacts with components of the ECM via its N-terminal multimerization domain. Moreover, COMP-Ang1 and CMP-Ang1 show a more robust Tie2 activation in Hela cells with high levels of ectopically expressed Tie2, hence suggesting a superior paracrine activity of COMP-Ang1 and CMP-Ang1 as compared to wt-Ang1. In fact, paracrine Tie2 activation was demonstrated only with COMP-Ang1 in HPMEC. COMP-Ang1 was shown to form solely pentamers, this being most likely due to the presence of the COMP multimerization domain. In contrast, CMP-Ang1 was shown to preferentially form dimers owing to the presence of its CMP dimerization domain **(Fig. 1D).** Tie2 activation has been previously reported to require a specific distance of the Ang1 binding domains in relation to the two Tie2 monomers (Leppänen et al. 2017). This very specific steric/spatial interaction is potentially facilitated by the pentameric COMP-Ang1 as compared to the dimeric CMP-Ang1 in low Tie2 expressing HPMEC cells. As a result, the observed higher and robust COMP-Ang1 mediated Tie2 activation in our system is most likely caused by its higher solubility compared to wt-Ang1, as well as by its more efficient multimerization into active pentamers as compared to that of CMP-Ang1. The difference in Tie2 activation **(Fig. 2)** in our hands led us focus on mRNA-76 encoding COMP-Ang1 for our further experiments. We subsequently demonstrated that mRNA-76 mediated COMP-Ang1 expression and Tie2 activation had phenotypic effects on cell adhesion by way of monitoring transcellular electrical resistance in HPMEC monolayers **(Fig. 3).** This observed stabilization of endothelial barrier function in *in vitro* cell culture is consistent with our *in vivo* results demonstrating a reduction of pneumolysin-evoked permeability in isolated perfused and ventilated mouse lungs after mRNA-76 treatment **(Fig. 6).** The underlying LNP formulation used in these experiments is based on our previous experimental work with siRNA-lipoplex formulations (Fehring et al. 2014; Santel et al. 2006b, 2006a). Functional delivery of the mRNA-76 payload was shown to occur in the lung vasculature, more specifically in pulmonary capillary endothelial cells (scRNA-Seq **Fig. 5).** Hence, here the synthetic mRNA becomes translated and consequently respective COMP-Ang1 is secreted as a multimer in order to bind and activate Tie2 receptors in a presumably paracrine and autocrine rather than endocrine manner **(Fig. 4).** Our data demonstrating COMP-Ang1 expression in mouse lung tissue together with *in vivo* mRNA-76 uptake by single cell transcriptional analysis and its subsequent Tie2 pathway activation in moues lung tissue therefore demonstrate the projected mode of action of our novel mRNA-76-cLNP compound. Data in **Fig. 9 B** show a significant decrease in LPS induced neutrophil influx into the alveoli after single mRNA-76-cLNP treatment. We did not observe changes in LPS induced vascular leakage (e.g., changes in wet lung weights) with mRNA-76-cLNP when compared to control mRNA formulation in this model. These data may indicate that vascular leakage and leucocyte extravasation do not necessarily occur together in blood vessel of the lung, which has been discussed previously by others (Filewod and Lee 2019). Generally, these findings in murine models for LPS-induced lung injury may also depend on the time point at which samples are obtained and the corresponding physiological data are captured. In contrast to the IPML model for primarily addressing vascular leakage, the LPS model might be not adequate for simultaneously addressing both neutrophil efflux and lung edema at 24 h post endotoxin challenge. Intravenous mRNA-76-cLNP treatment was shown to intervene with leukocyte and in particular neutrophil extravasation into the alveoli after LPS challenge. It should be noted however, that the reduction of the influx of leukocytes in addition to a reduction in edema formation might be clinically highly desirable to counteract the delayed apoptosis of parenchymal neutrophils believed to contribute to tissue injury (Teder et al. 2002).

Taken together the data in our IPML mouse model demonstrate a dose dependent therapeutic effect of mRNA-76-cLNP regarding endothelial permeability, while the LPS-induced pulmonary neutrophilia mouse model depicted in **Fig. 9.** shows therapeutic efficacy on dysregulated lung inflammation. Both underlying pathophysiological mechanisms, namely loss of endothelial permeability and dysregulated lung inflammation, are hallmark events at the onset of early/ mild ARDS. Thus, the advantage of our novel mRNA-76-cLNP compound as described above is its ability to act in a highly spatially restricted manner thereby enabling the Tie2 agonist to rapidly and almost exclusively target specifically that site of the endothelial capillary bed where the pathophysiological lung edema occurs. This particular strong spatial restriction of action of our mRNA compound enables a more rapid onset of Tie2 pathway activation, as well as improved pharmacodynamics, compared to those drug modalities which require a more continuous treatment due to the rapid clearance of recombinant proteins from the systemic circulation. Increased Ang2 levels were also found more recently to be strongly associated with COVID-19 in-hospital mortality and with non-resolving pulmonary condition (Villa et al. 2021a). Thus, Ang2 may be an early and useful predictor and biomarker for patient stratification and of the clinical course of COVID-19, and impending Ang2 binding a potential treatment option against COVID-19 (Villa et al. 2021b). Moreover, inhibition of the antagonistic Ang2 or a transient and localized upregulation of Ang1 -or a hyperactive derivative thereof- is a promising strategy to restore Tie2 signaling in order to stabilize the barrier function and to prevent the inflammatory lung leakage in ARDS/COVID-19 patients. With the understanding that the angiopoietin/Tie (Ang/Tie) family has an established role in vascular physiology in regulating angiogenesis, vascular permeability, and inflammatory responses, our data corroborate effectiveness and mode of action of the clinical approach of treating damage of the lung vascular epithelium with transient expression of an mRNA-LNP modality. The highly specific targeting of our respective COMP-Ang1 cationic LNP of the lung vasculature thus enables spatially highly restricted expression of COMP-Ang-1 for treating edema formation in ARDS patients as a new treatment modality for further development.

